# Effect of perchlorate on biocementation capable bacteria and Martian bricks

**DOI:** 10.1101/2025.02.24.639804

**Authors:** Swati Dubey, Shubhanshu Shukla, Nitin Gupta, Rashmi Dixit, Punyasloke Bhadury, Aloke Kumar

## Abstract

With the recent discovery of perchlorate (0.5-1%) in Martian regolith, more experiments related to the impact of perchlorate on microbial life are crucial to understanding the possibility of earth life forms that could sustain on the Martian terrain. While we are familiar with the idea of bioconsolidated Martian bricks made via Microbially Induced Calcite Precipitation (MICP), studies on the effect of perchlorate on Martian bricks & biocementation capable microbes have been obscure. In this work, we investigated the effect of perchlorate (MgClO_4-_ salt) on a lab-isolated biocementation capable bacteria & Martian bricks bioconsolidated by the same, with 1% perchlorate in Mars Global Simulant-1 (MGS-1). The screening of biocementation-capable bacteria involved phenol red assay for urease activity followed by Scanning Electron Microscopy (SEM) and X-ray diffraction (XRD) study of the precipitate formed through MICP via ureolytic pathway. The biocementation capable bacterium SI_IISc_isolate was found phylogenetically closest to *Sporosarcina pasteurii* strain S2135, its draft genome being approximately 3.69 Mb. To understand the effect of perchlorate on SI_IISc_isolate, we majorly relied on Gram-staining & SEM. The negative effect of perchlorate stress on the isolate was evident by its decreased growth in the presence of varying concentrations of perchlorate through plate assays, growth curve studies in broth & live-dead staining. Gram-staining study and SEM both revealed that perchlorate induces the release of extracellular matrix (ECM) and promotes clustering of cells by the bacteria, which we termed as ‘multicellularity-like behavior.’ Further, we constructed Martian bricks with Martian Global Simulant (MGS-1) along with 1% perchlorate, utilizing the microbially induced calcite precipitation ability of the *Sporosarcina* sp. strain SI_IISc_isolate via ureolysis, following an established protocol at our lab. The bioconsolidation experiments showed that in the presence of a natural adhesive – guar gum, perchlorate tends to significantly improve the compressive strength of Martian bricks. However, the end result eventually relies on the overall effect of various additives in the regolith.

## I. INTRODUCTION

Ancient Mars has had a history of water^1–3^ and the present Mars has an uncanny resemblance to the early atmosphere of Earth ^4^; the red planet is an interesting study into planetary evolution as well as into the origin and evolution of life.^5–7^ The red planet has also been identified as a potential candidate for the establishment of a human colony; our intended progression from manned missions into space (Low Earth Orbits) to Mars Missions by 2030, demands extensive research on Mars habitation with a special focus on sustainable and cost-effective methods. In-Situ Resource Utilization (ISRU) is an idea that encourages the utilization of locally available resources for space exploration.^8^ An abundant resource on Mars is its regolith^9,10^. The Martian regolith is mostly composed of pyroxenes, olivines, plagioclase, and abundance of iron oxides.^11,12^ Along with these minerals, the perchlorates were detected in 2008 by the Phoenix lander^13,14^ – a discovery that was confirmed by the instruments on board the Curiosity rover.^15,16^ Figure 1 represents the Martian landscape where the landing sites of the rovers and perchlorate detection are delineated.

**Figure 1.**
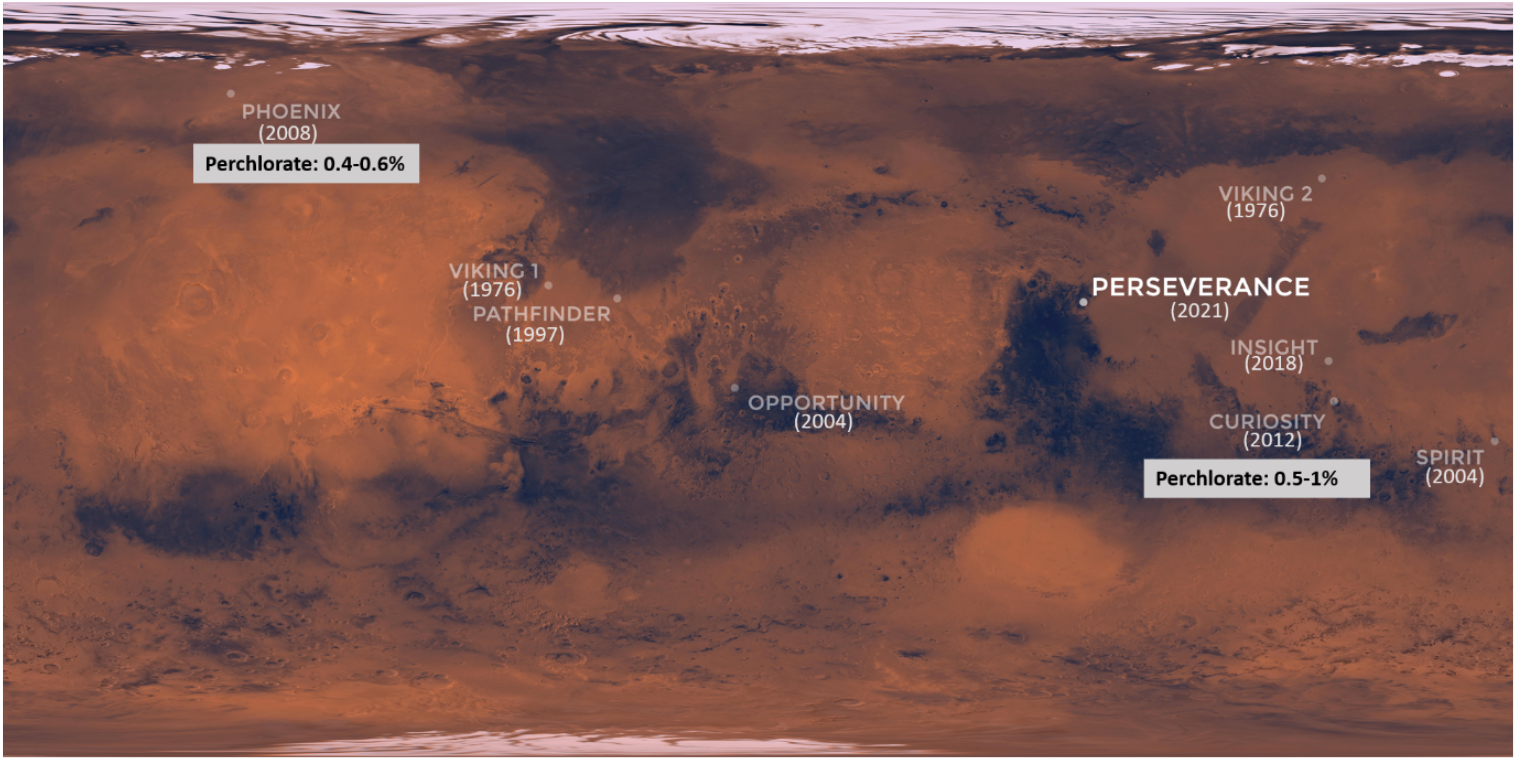
Modified map showing the landing sites for landers sent by NASA for Mars exploration. The respective landing years are mentioned in brackets. The Mars visit which found perchlorate on the Martian surface is highlighted. (Image Credits: NASA-JPL/Caltech)

**Figure 2.**
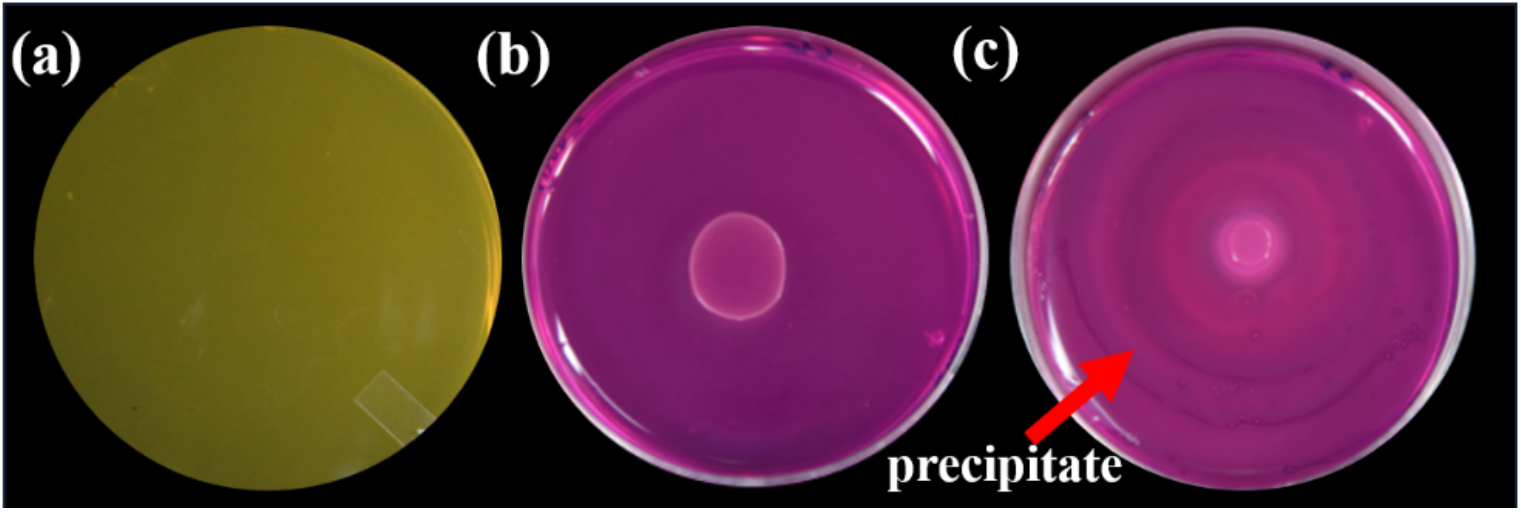
Phenol Red plate Assay for SI_IISc_isolate (also called SI) to test for ureolysis activity. (a) represents Control_ No bacteria on SMU+ 0.001% Phenol Red Dye. (b) SI spot inoculated on SMU+ Dye. The colour change from yellow to pink represents the pH change of the media from ∼5.5 to .8 and above. (c) represents SI spot inoculated on SMU+ Dye + 25mM CaCl2. The white lines on the plate represent calcite precipitate. Note: Plate dimensions-90×14mm.

Research work that needs to employ Martian regolith typically employs ‘simulants’, which simulate closely the content of actual Martian regolith. Earth-based experiments using Martian regolith simulant are an important step towards understanding how Earth-based life can thrive on Mars. For example, the BIOMEX experiment used Martian regolith simulant (MRS) as a substrate to test the viability and growth of *Deinococcus radiodurans* and other microbes on the International Space Station (ISS).^17^ Martian bricks have been constructed with Mars Global Simulant-1 (MGS-1) by using a biocementation-capable bacteria *Sporosarcina pasteurii*.^*18,19*^. *S. pasteurii* is a well-known bacterium for its biocementation capability through a mechanism called microbially induced calcite precipitation (MICP) via ureolysis. ^20^ Harris et al. demonstrated soil fertility by studying *Sinorhizobium meliloti*-legume interactions in JSC Mars 1 simulant.^21^ However, Earth-based simulants like MGS-1, do not have perchlorate that is present in the actual Martian environment.

Despite the fact that perchlorates are a minority constituent of Martian regolith they are relevant because they have been closely associated with the habitability of the planet. Being a strong oxidant, they can readily degrade organic matter and hence have been claimed to interfere with the detection of biosignatures in the regolith. Since the hydrated perchlorates are seen as sinks for H_2_O and have also been shown to inhibit the salt and ice crystallization,^22–26^ their study in context to regolith and microbial life could yield interesting insights. On Earth, perchlorates are identified as an environmental pollutant, potentially toxic to living cells by causing severe oxidative stress.^27–29^ However, certain perchlorate metabolizing bacteria belonging to genera such as *Sedimenticola, Marinobacter, Azoarcus, Dinitromonas*, and *Marinobacter*^30^ and perchlorate tolerant bacteria such as *Haloterrigena* sp. strain SGH1^31^, *Halomonas salifodinae*^*32,33*^ are known to us. Most of these microbes are extremophiles-a group that has been most frequently associated with the study of extraterrestrial possibility of life.

Biocementation-capable microbes as our tools of a sustainable Mars habitation through their ability to consolidate the Martian regolith along with their potential to withstand several harsh conditions^34–39^, are good candidates for examining the regolith-microbe interaction and are rarely studied with this perspective.^40–42^

In our work, we investigated the impact of perchlorate on *Sporosarcina spp*. strain SI_IISc_isolate (hereafter interchangeably called SI) isolated by us. The strain’s biocementation capability is identified through biochemical assays, scanning electron microscopy, and x-ray diffraction (XRD) study. 16S rRNA sequencing and whole genome sequencing were done for phylogenetic identification and to identify the metabolic potential of the bacteria. The Minimum Inhibitory Concentration (MIC) of perchlorate for the SI was identified through plate assays and growth curve study, after which the impact of 0.5%, 1%, and 2% perchlorate on the bacteria was studied through optical microscopy after gram-staining and through scanning electron microscopy (SEM). Furthermore, the biocementation ability of SI was explored through the construction of Martian bricks by a previously established protocol using MGS-1 simulant with the addition of 1% perchlorate.

## II. MATERIALS & METHODS

### Screening of MICP-capable bacteria

The soil sample was collected from a site in Bangalore, India during March 2021. The collected soil sample was immediately incubated at 32^0^C in Synthetic Media + 2% Urea solution for 24 hrs and microbial isolation was done according to the serial dilution method and subsequently, colonies of distinct different morphologies were selected for further screening. The isolates were subjected to biochemical tests including for carbohydrate utilization using Bacillus Identification Kit (HiMedia, India). Gram Staining was done to study the cellular morphology.

### Screening for Urease activity

A qualitative phenol red assay was used to determine the ureolytic activity of the isolated strains. Synthetic Media+ 2% Urea (hereby called SMU) along with 0.001% phenol red dye (pH indicator) was used for the assay. Spot inoculation of different colonies was done on the plates followed by incubation for 18-24 hrs at 32^0^C. Based on positive results for phenol red assays, bacterial strains were subsequently selected to test for their microbially induced calcite precipitation or MICP ability through the formation of CaCO_3_ precipitates, via ureolysis.

### Microbially Induced Calcite Precipitation (MICP)

For screening the bacterial isolates for their ability to form CaCO_3_ precipitates in-vitro, 25mM of Calcium Chloride (diluted from 1M of CaCl_2_ stock solution) was added to synthetic media with 2% urea (SMU) broth. This media was inoculated with the selected bacterial strains in triplicates and left for incubation at 32^°^C for 5 days on a shaker incubator (BioBee, India), at a rotating speed of 130 rpm, along with control tubes with no bacterial inoculation.

### Scanning Electron Microscopy of the precipitate

Based on the MICP screening, an isolate -referred to as SI_IISc_isolate or SI, was selected. The total precipitate formed by this bacteria was subjected to scanning electron microscopy (SEM) to study the morphology of the precipitate formed in the presence of calcium and urea, by the ureolytic activity of the bacteria, The total precipitate including the cell biomass was therefore fixed with 2.5%glutaraldehyde: 2%formaldehyde PBS buffer (pH 7.4) in the ratio of 1:1:4 for at least 30 min, followed by centrifugation at 5^°^ C and 5000 rpm for 5 minutes followed by application of standard steps of SEM. Before SEM imaging, dried samples were coated with gold in a sputter coater. Samples were observed under a SEM microscope (Carl Zeiss AG— ULTRA 55, Germany).

### X-Ray Diffraction study of the precipitate

To identify the mineral phase of the precipitate formed by SI, an XRD study was done. The precipitate was pelleted down by centrifugation (5000rpm, 10min) and then air dried, followed by grinding by mortar and pestle for homogeneity. This precipitate was further analyzed for Powder-XRD study through an X-Ray Diffractometer (PANalytical Philips diffractometer) using CuKα Rays (λ = 1.54056 A), diffraction angle 2θ ranging from 5^0^ to 90^0^. The raw data was analyzed through X’Pert Highscore Plus Pattern Analysis Software using ICSD (Inorganic Crystal Structure Database) sub-files.

### Genomic DNA extraction of SI_IISc_isolate

Bacterial cells were scraped out from the agar plates and suspended in 1ml of 1X TE (Tris-EDTA, Tris-Cl 10 mM pH 8.0, EDTA 1 mM) before undertaking extraction of genomic DNA. Finally, the pellet was air-dried and dissolved in 1X-TE buffer.

### PCR amplification, sequencing of 16S rRNA fragment, and molecular phylogeny

For PCR amplification of the 16S rRNA of SI, two primers 203F and 1492R were used^43^. The PCR reactions were set following the protocol as described earlier. The PCR amplicon was purified and subjected to Sanger sequencing in the ABI 3130xl platform. Based on the blastn result, the top 10 sequences showing hit with the sequence data of SI_IISc isolate were considered. The sequences were aligned using BioEdit and subsequently, the alignment was subject to ModelTest in MEGA 11. Based on the output of ModelTest, the Maximum Likelihood (ML) tree was constructed with a bootstrap support of 1000. The sequence of the SI_IISc isolate has been submitted to GenBank and the accession number is PP919058.

### Whole genome sequencing of SI_IISc isolate

High-quality genomic DNA from SI was extracted using a modified protocol. The whole genome was sequenced on a MinION platform using Oxford Nanopore Technologies (ONT) Sequencing chemistry. The genome library was prepared using a ligation sequencing kit (SQK-LSK109, ONT, UK). The library was purified using magnetic beads (Sergei Lab Supplies LLC, USA) and sequenced on an R9.4.1 Flowcell. The *de novo* genome assembly was filtered for long reads using flye (v2.9.3)^44^. The quality of the assembled genome was evaluated using QUAST (v5.2.0) and annotated using Prokka (v1.14.6)^45^. The genome map was created using Proksee^46^.

### Effect of perchlorate on SI_IISc_isolate

To understand the effect of perchlorate on the growth of the *Sporosarcina sp*. strain SI_IISc_isolate including inhibition, the minimum inhibitory concentration (MIC) of perchlorate was determined using plate assay and broth tests. In both broth and solid media, perchlorate (as magnesium perchlorate) was introduced in the form of a solution in the Synthetic Media with 2% Urea or SMU, as described earlier. To study the effect of perchlorate on SI_IISc isolate at cellular & sub-cellular levels, Gram-staining and SEM were performed in the presence of 0.5%, 1%, and 2% of perchlorate in SMU & Nutrient Broth (HiMedia, India) +2% Urea (NBU) broth media respectively. For gram-stained bacterial cells, imaging was undertaken at 100x magnification under a bright field microscope. (Leica DMi8, USA). Similarly, SEM images of the bacterial cells were undertaken, following the protocol as described earlier. Moreover, to understand the effect of perchlorate on the viability of SI, live/dead staining was performed (refer to Supplementary file S1).

### MICP consolidated Martian bricks with perchlorate

The Martian bricks were constructed utilizing the biocementation ability of the *Sporosarcina sp*. strain SI_IISc_isolate using MGS-1 as Martian regolith simulant along with 1% perchlorate by the protocol previously established at our lab^47^. To understand the role of different components of the slurry mixture used for our brick consolidation experiment, various controls were used (discussed later in the results section). The cubical brick samples obtained from this method were uniformly ground and polished using a portable grinder (Bosch GWS 600) to ensure consistent compression behaviour. The final specimens were in the form of cubes with dimensions of 19±1 mm. Finally, the compression strengths of the bricks were evaluated using a Universal Testing Machine (Instron-5697) equipped with a 5 kN capacity load cell and operated at a 0.5 mm/min loading rate.

## III. RESULTS AND DISCUSSION

### A. Isolation of MICP-capable bacteria

#### Microbially Induced Calcite Precipitation

Only one isolate named SI_IISc_isolate gave positive results for urease activity through phenol red assay. The isolate formed amorphous precipitates in the presence of 25mM CaCl_2_ in Synthetic Media+2% Urea (SMU), after 5 days of incubation (Fig.2). Similar qualitative analysis of ureolytic activity (phenol red plate assay) was performed with different concentrations of perchlorate to understand the impact of perchlorate on the ureolytic activity of bacteria (Fig. S2).SI formed a precipitate in the presence of 25mM CaCl_2_ in SMU broth after 5 days of incubation at 32^0^ C. The total precipitate hence formed was pelleted down after centrifugation and studied through SEM and X-Ray Diffraction to validate the MICP ability of SI bacteria.

The precipitate formed due to ureolytic activity by SI when observed through SEM showed floret-like precipitates along with rhombohedral nanocrystalline structures which is a characteristic morphology for calcite precipitates. The bacterial cells were also seen wrapped up to the floret-like precipitates as shown in figure 3(a).

**Figure 3.**
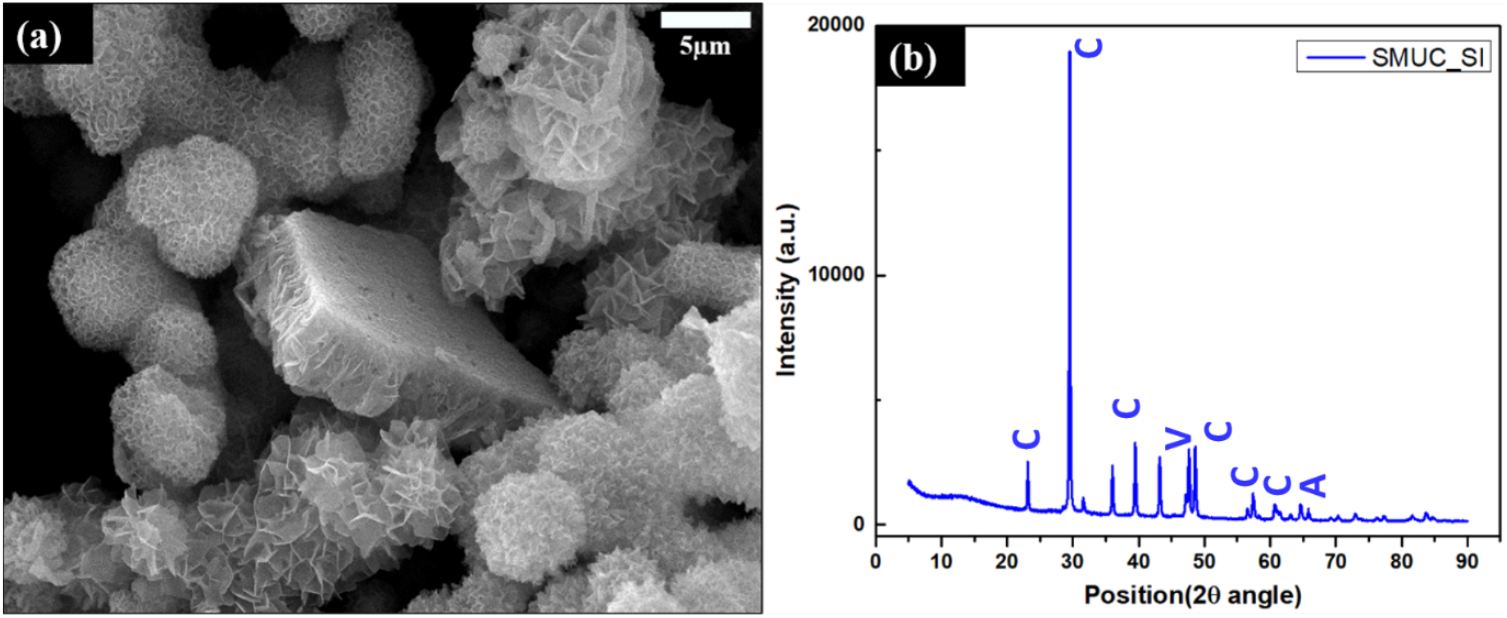
(a) Scanning Electron Microscopy images of the calcite precipitate made by the MICP-capable bacteria SI via its ureolytic activity. (b) X-Ray Diffraction (XRD) results for the precipitate. C represents Calcite, V represents Vaterite, A for aragonite, and peaks as identified through the ICSD database sub-file.

Analysis of the XRD pattern of the precipitate revealed the presence of calcite peaks along with a few peaks of vaterite and aragonite, as analyzed through X’pert HighScore Plus Pattern Software. Identification of mineral precipitates was done through thoroughly indexed peaks obtained at different 2θ & hkl values along with d-spacing, studied in reference to the available database from the ICSD sub-files. Figure 3(b) is the graph for the relative intensity at different diffraction angles (2θ), which shows calcite peaks, as identified through thoroughly indexed peaks, for which the intensity values, diffraction angles along with the relevant reference IDs are given in table S2.

#### Molecular identification and phylogeny of SI_IISc isolate

A gram-positive, facultatively anaerobic bacterial isolate SI_IISc_isolate (bacterial morphology can be seen in the inset of Fig. 4) gave positive results for phenol red assay and was able to cause calcium carbonate precipitation in-vitro, through its ureolytic activity. The summary of biochemical test results along with the motility test for the isolate has been given in table S1 & figure S1 respectively in file S1. Based on blastn analysis, the 16S rRNA sequence of SI_IISc isolate showed 99% identity with *Sporosarcina pasteurii* strain S2135 (Accession Number: 718761), a bacterium that was isolated from the soil environment in China as well as with *Sporosarcina pasteurii* strain WJ-4 (Accession number KC211296), a bacterium isolated from the concrete environment in Korea. Based on ML phylogeny, SI_IISc_isolate clustered with the *Sporosarcina* clade with strong bootstrap support but also indicated that the isolate from this present study could be potentially novel.

**Figure 4.**
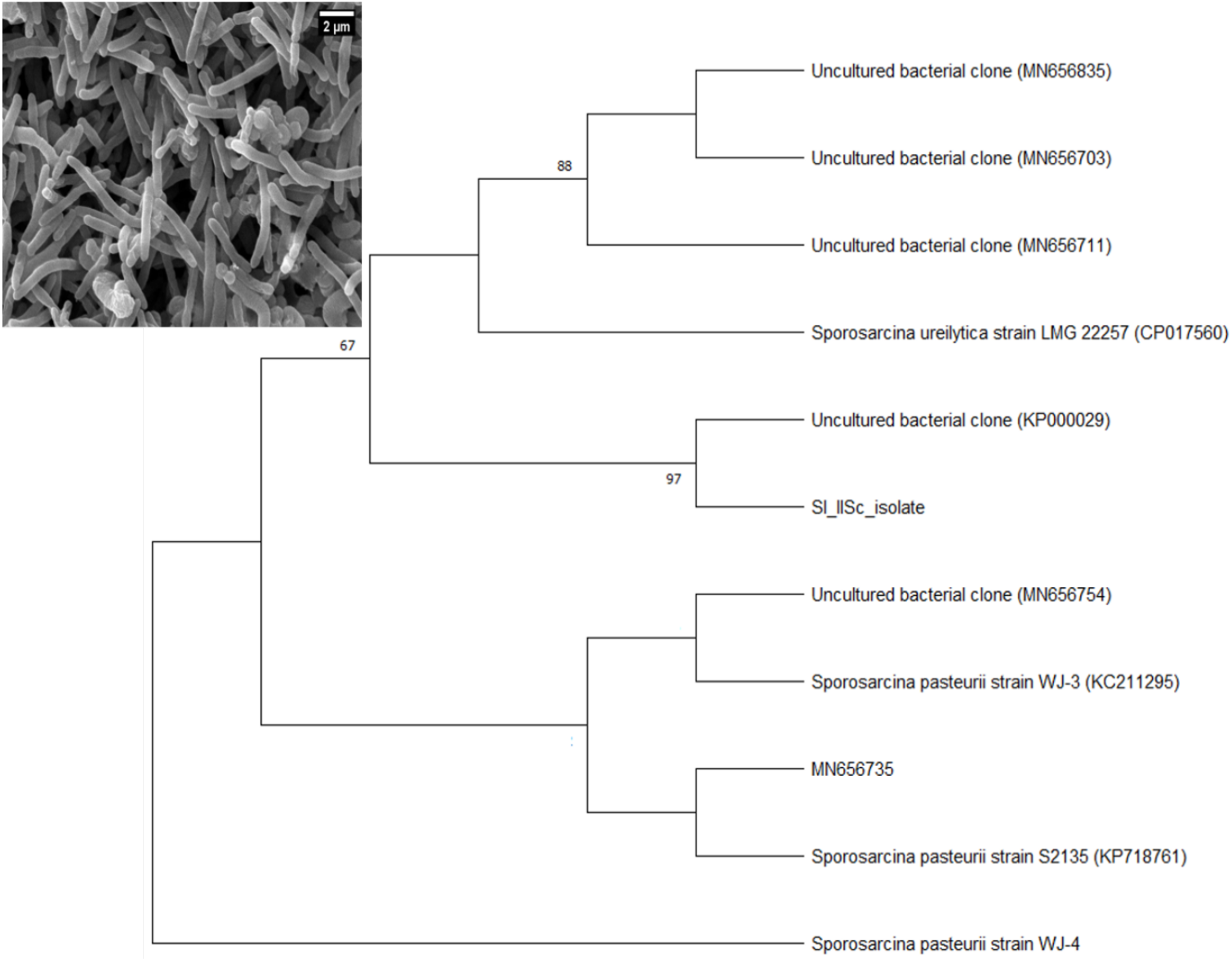
Maximum Likelihood tree of 16S rRNA showing the relationship of SI_IISc isolate with closely related taxa, values in node indicate bootstrap support. Inset is a SEM image of the SI_IISc isolate.

The genome size of *Sporosarcina* sp. strain SI_IISc_isolate is approximately 3.69 Mb in size. The draft genome comprises 27 ribosomal RNAs, 81 transfer RNAs, 1 transfer messenger RNA (tmRNA), and 4,678 coding sequences (CDS). In total 4678 CDS have been identified from the draft genome. The genome map is shown in figure 5. The identified genes (4787) code for many metabolic pathways including for nitrate reductase that play an important role in the utilization of perchlorate. The UreC gene attributing to ureolytic activity of the bacteria has been also identified from the draft genome, which is available in NCBI database under the BioProject ID: PRJNA1221533 and the BioSample number: SAMN46743258.

**Figure 5.**
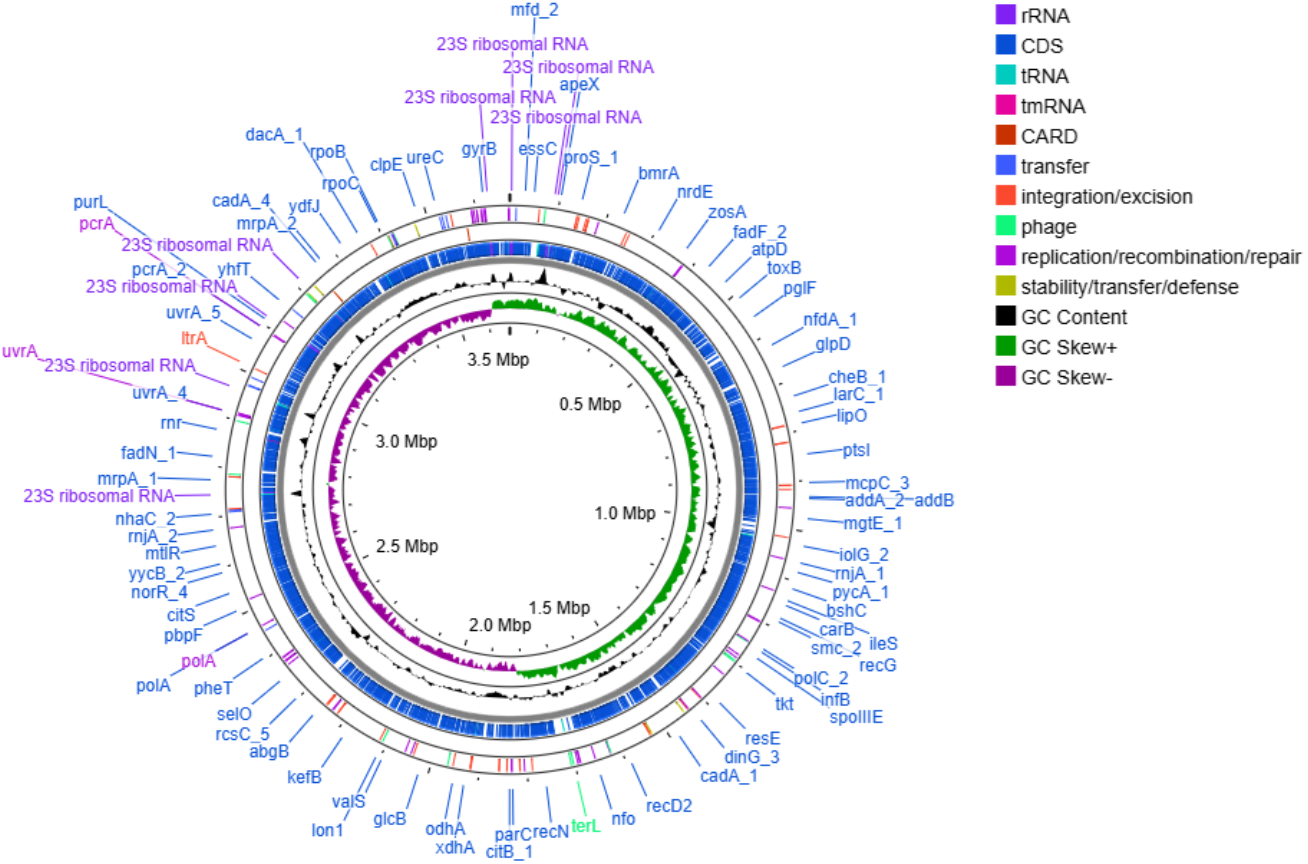
Genome map showing a circular map of the genome (representation only), identified genes, GC Content, and GC Skew of SI_IISc isolate

## B. Effect of Perchlorate on SI_IISc_isolate

To know the effect of perchlorate on the growth and viability of the *Sporosarcina sp*. strain SI_IISc_isolate, we performed a growth curve in different concentrations of perchlorate, and the Minimum Inhibitory Concentration (MIC) of perchlorate on this strain was determined to be 3%, through plate and broth assays exhibiting tolerance up to 2% of perchlorate salt. The growth curve experiments showed a gradual decrease in growth of this strain with increasing concentration of perchlorate (refer Fig. S3 & Fig. S4). The negative impact of perchlorate on this strain was also validated through live-dead staining, performed in NBU media in 0.5%, 1%, and 2% concentrations of perchlorate salt along with a control (no perchlorate) set. After 8 hours of incubation, a gradual increase in the number of dead cells was observed with increasing concentrations of perchlorate salt and the bacterial cell size also diminished under perchlorate’s presence (as compared to the control) (refer Fig. S5)

Based on these results which confirmed that perchlorate was a stressor for this strain; to further study the impact of perchlorate on the biocementation capable bacteria we performed Gram-staining and SEM studies of this isolate in the presence of 0.5%, 1%, and 2% perchlorate. Since perchlorate found in Martian regolith is known to be in the range of 0.5%-1%, we performed our experiments with 0.5%, and 1% along with 2% perchlorate salt concentration with 2% perchlorate concentration as an outlier.

### Gram staining

On gram staining the bacterial cells being exposed to different concentrations of perchlorate, we found certain changes in the morphology of the individual cells along with changes in their cells-distribution pattern. Figure 6 (a) to (d) shows gram-stained images with their corresponding magnified versions in the insets attached below. Since SI being phylogenetically assigned to *Sporosarcina sp*., is a gram-positive bacterium; the bacterial cells were stained purple, elongated rod-shaped cells as shown in figure 6a.

**Figure 6.**
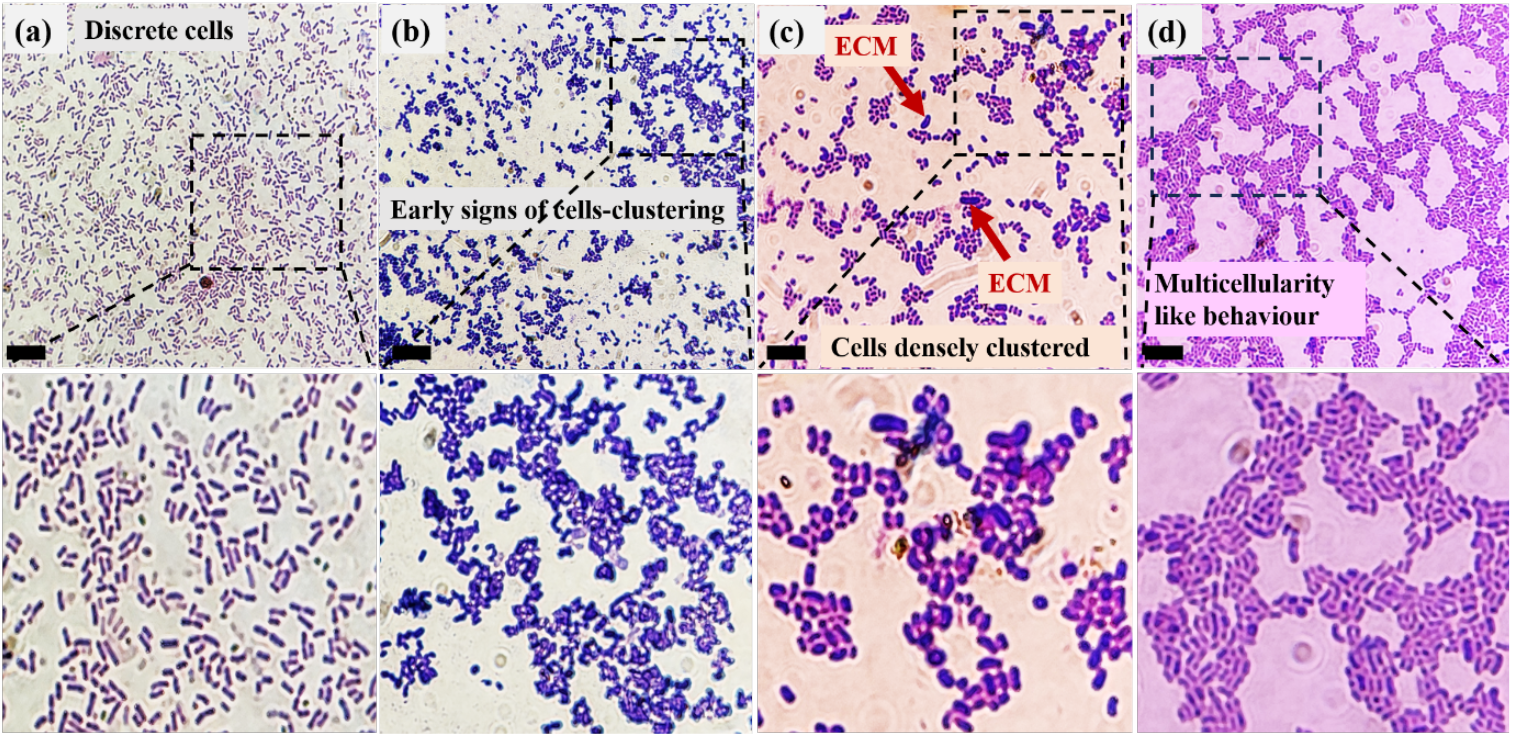
Gram-staining results showing the difference in cell morphology and behaviour of the *Sporosarcina sp. strain* SI_IISc _isolate(or SI) (a) represents SI Control (b) SI + 0.5%Perchlorate (c) SI + 1%Perchlorate (d) SI + 2%Perchlorate. The lower panel represents magnified portions of corresponding images in the top panel. The scale bar represents 10µm.

On multiple repetitions of the gram-staining experiment, we found that the *Sporosarcina sp*. strain SI_IISc_isolate cells exposed to perchlorate tend to adapt a smaller size & less elongated shape than the control set. A more pronounced change was in the distribution pattern of the cells in response to perchlorate. It was noted that on increasing the concentration of perchlorate, the tendency of bacterial cells to form clusters gradually increased. For example, the bacterial cells exposed to 0.5% perchlorate, showed early signs of clustering by forming small undefined clusters in figure 6b.as compared to the control in figure 6a., the cells with 1% perchlorate exposure in figure 6c., formed very small groups of cells in a better-defined cluster, while the cells in figure 6d. with 2% perchlorate exposure showed very well-defined clusters formed more repetitively than the other two sets. However, it would be early to comment on whether there was a particular pattern in the clustering which led us to believe that perchlorate’s presence encouraged multicellularity-like behaviour in the bacteria. We also noticed that the perchlorate-exposed cells released extracellular matrix (ECM) seen as purple patches or blots on the slide and this response was observed in cells exposed to all three concentrations (0.5%, 1%, 2%) of perchlorate, more clearly seen in figure 6c (indicated by red arrows). This observation was consistent with multiple repeats of the experiment which strongly indicates that perchlorate induced ECM release in cells.

### Scanning Electron Microscopy

For SEM analysis, *Sporosarcina sp*. strain SI_IISc_isolate cells were exposed to 0.5%, 1%, and 2% perchlorate (MgClO_4-_ introduced as salt solution) in SMU broth followed by 5 days of incubation after which the cells were centrifuged to obtain a pellet, fixed for SEM study. Figure 7. (a) to (c), (d) to (f) & (g) to (i) shows SEM images of the total pellet obtained after exposure of this strain to 0.5%, 1%, and 2% perchlorate respectively.

**Figure 7.**
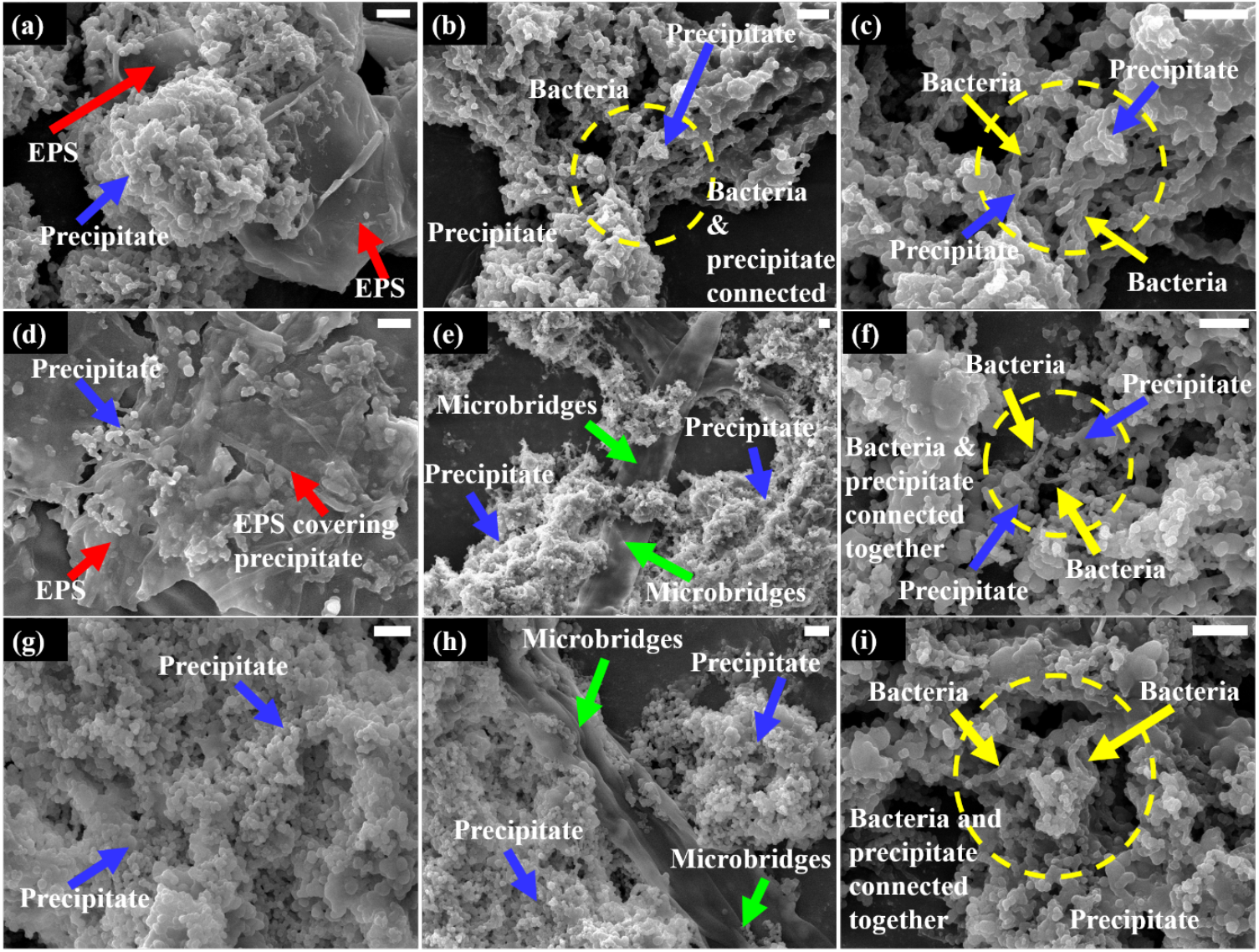
Scanning Electron Microscopy: Images of the total precipitate formed from bacterial exposure to different concentrations of perchlorate (as (MgClO_4_)_2_). Figures (a) to (c) represent SEM images for total precipitate with SI+ 0.5%Perchlorate (d) to (f) with SI+ 1%Perchlorate & (g) to (i) for SI+2%Perchlorate. The scale bar represents 5 µm.

It was observed during & after the incubation period that this strain induced the formation of an inorganic precipitate in the presence of perchlorate which dominated the total pellet. These precipitates can be abundantly seen in the SEM images indicated by blue arrows (morphology most clearly evident in figure 7g). The chemical nature of this inorganic precipitate could not be identified through XRD due to the pellet’s hygroscopic nature and impurity with high organic mass. However, a significant release of extracellular matrix (ECM) also referred to as extrapolymeric substances or EPS in the SEM images can be seen as organic mass more evident in figure 7a and 7d, indicated by red arrows. No ECM was observed in SEM images of the control – that is bacteria without perchlorate exposure (refer inset image in Figure 4). The release of ECM was an observation consistent with the gram-staining study and therefore it further validates that perchlorate encouraged the ECM release by this strain. Of special interest were the microbridges – a term we refer to the ECM material acting as a conduit between the salt precipitates and the bacteria, as shown in figure 7e and 7h, indicated by green arrows. The interaction of the inorganic precipitate and bacteria (indicated by yellow arrows) can be easily seen in images 7.b, c, f & i, indicated by yellow dotted circles. With increasing concentration of perchlorate, the occurrence of bacterial cells decreased along with significant reduction in ECM release, which could be attributed to the negative impact of perchlorate on the viability of bacterial cells. The response of the strain towards perchlorate in the form of ECM release, the occurrence of microbridges, and the interaction of bacterial cells & precipitate were seen to be more robust in another experiment (data not shown) with ‘conditioned’ bacterial cells that were pre-treated with 0.25% of perchlorate before repeating the experimental set-up as described in the paper(with ‘non-conditioned’ bacterial cells).

### C. MICP-consolidated ‘Martian bricks’ with perchlorate

#### Compressive Strength Evaluation

For bioconsolidation of bricks, a blend of the *Sporosarcina sp*. strain SI_IISc culture along with Martian regolith was combined with SMU media in a slurry form and allowed to incubate within the mold cavities for approximately 5 days. Various control experiments were set up simultaneously such as the slurry without perchlorate, guar gum, and nickel chloride respectively, and in different combinations in order to understand the effect of perchlorate on bioconsolidation. The general workflow of the bricks-casting method can be understood by the schematics in figure 8. Following the incubation period, the resulting cubical samples were extracted, dried, and subjected to quasi-static compression tests.

**Figure 8.**
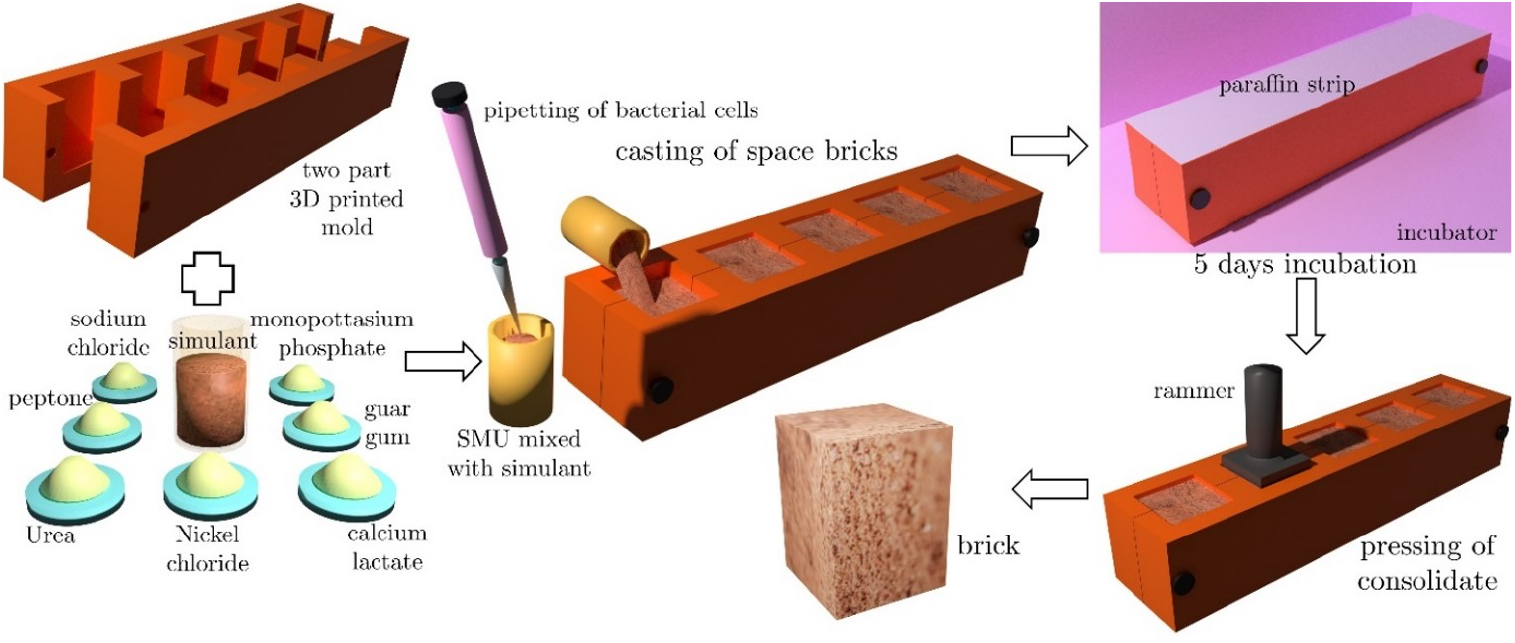
Schematic depicting the brick casting procedure alongside the consolidation process facilitated by MICP in simulated regolith.

The results of compressive strength measurements derived from different slurries with various media combinations are given in figure 9. The Martian bricks made with MGS-1 consolidated with deionized (DI) water yielded a very low compressive strength of 0.17MPa and crumbled under self-weight. The bioconsolidation with the culture resulted in a strength comparable to the corresponding abiotic control and the slurry mixture including bacteria along with 1% perchlorate (denoted by SI+P) also yielded similar results with bricks’ compressive strength being 0.064 MPa. These results can be attributed to the polyextreme conditions in the Martian regolith, especially in the presence of perchlorate. However, using 1% guar gum which is a natural adhesive ^51^, increased the compressive strength of bricks to 0.82MPa in the abiotic control (denoted by DI+G) and to 3.07MPa in the regolith mixture with bacteria (SI+G). This highlights the crucial role of guar gum in significantly encouraging the consolidation process and also confirms the contribution of this strain towards bioconsolidation aided by guar gum. The nickel combined with this strain and 1% perchlorate (SI+N+P) showed a ten-fold increase in the compressive strength, it being 0.64MPa, as compared to the corresponding control without nickel (SI+P). The compressive strength of bacteria with guar gum and nickel (SI+G+N) was 4.4MPa as compared to the without nickel mixture (SI+G, *σ*_*c*_ being 3.07MPa). These observations validated the role of nickel, a known catalyst for ureolysis ^52^, towards improved bioconsolidation via ureolysis-induced MICP.

**Figure 9.**
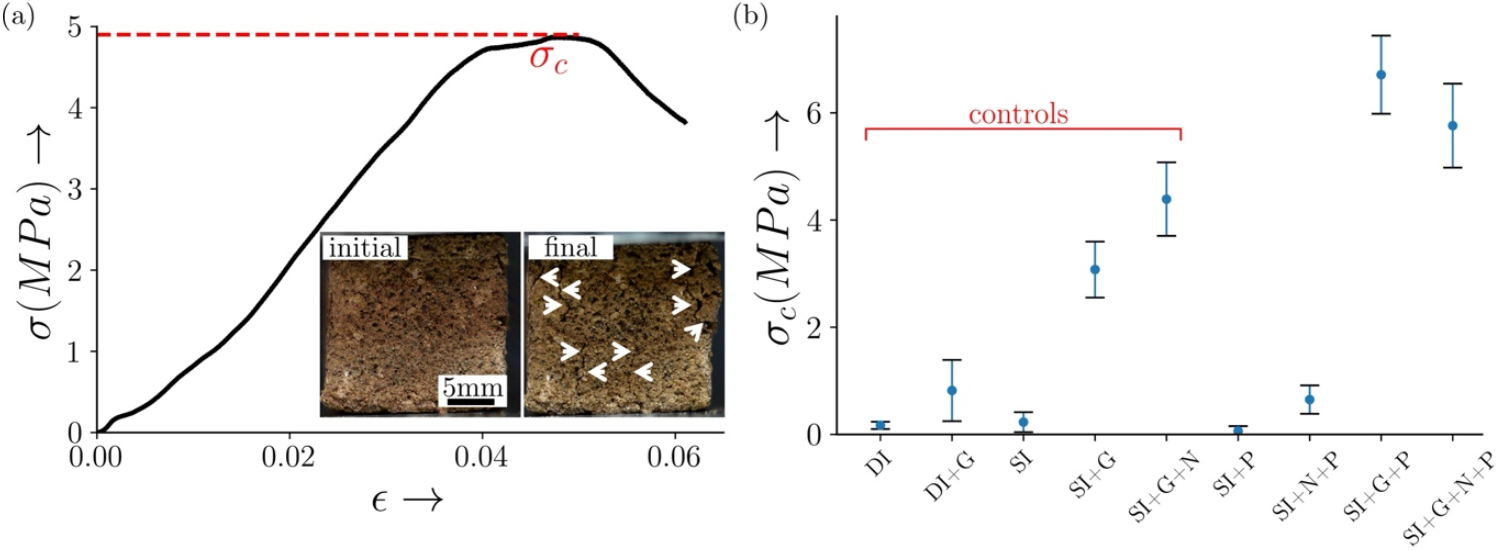
Panel (a) illustrates the stress-strain behaviour of the bio-consolidated specimen subjected to compressive loading. The peak stress observed was designated as the compressive strength σ_*c*_ of the consolidated brick. The inset provides a visual representation of the Martian brick specimen ‘s (made with Martian regolith+ SI+G+N+P) condition during loading, displaying the initial state of the specimen on the left and its final state on the right, with arrows indicating the locations of cracks. Panel (b) presents the compressive strength σ_*c*_, averaged from a minimum of four specimens across various treatment conditions. Error bars indicate the variability in brick strength. The treatments DI and DI+G were treated as a control to check for the significance of MICP and compared with SI and SI+G, respectively. The treatments examined include SI, SI+G, and SI+G+N, serving as controls to assess the influence of perchlorate on the compressive strength of the bricks, a control to check for the significance of MICP and compared with SI and SI+G, respectively. The treatments examined include SI, SI+G, and SI+G+N, serving as controls to assess the influence of perchlorate on the compressive strength of the bricks, a measure indicative of MICP mediation.

On the other hand, the compressive strength of bacteria with guar gum including 1% perchlorate (SI+G+P) was noted to be 6.71MPa, which was almost twice the compressive strength of the corresponding bricks without perchlorate control (SI+G). The bricks made with the bacteria, perchlorate, guar gum and nickel (SI+G+N+P) also showed higher compressive strength being 5.77MPa as compared to the corresponding without perchlorate set (SI+G+N) with *σ*_*c*_ being 4.4MPa. These results indicate that perchlorate enhanced the biocementation process in the presence of bacteria, where the role of guar gum was critical, acting as an adhesive and possibly a source of nutrition helping bacteria to survive the Martian regolith’s high salt conditions. However, comparing the SI+G+N+P sample as compared to the without perchlorate set (SI+G+N) showed a little decrease in the bricks’ compressive strength which can be attributed to the high metals/ ions stress to the bacteria due to the presence of nickel and perchlorate together, making bacterial survivability a bit challenging.

To conclude, these experiments showed that in the presence of 1% guar gum-an important binding agent; the strain showed efficient biocementation via ureolysis-induced MICP, improved further in the presence of 1% perchlorate. The role of nickel in improving biocementation was also identified. Overall, these findings underscore the complexity of MICP-mediated consolidation and the significance of understanding the interplay between various additives for optimizing strength in the Martian regolith-based constructions.

## IV. DISCUSSION

The bacterial extracellular matrix with or without biofilm in particular is known to encourage cells migration^53^, genetic exchange, capturing of ions, cell signalling. & has been frequently correlated with environmental stress as a means of protecting the cells against antibiotics, heavy metals stress, oxidative stress or to overcome the lack of nutrients etc. ^54–57^

In an experiment with *Bacillus cereus*, high calcium concentration acted as a stress for the bacteria in response to which, the bacteria released ECM^58^. Since our experiments clearly indicate that perchlorate acts as a stressor for the *Sporosarcina sp*. strain SI_IISc, the release of ECM is likely a coping mechanism that can be validated with further studies. In the same experiment with *B. cereus*, ECM has been seen as a link between biofilm and the mineralization ability of the bacteria. However, in our case, more experiments would be required to examine that. In another study with *S*.*aureus* biofilms, their ECM material was seen as a means of connecting the microbes together suggesting a communication between the multicellular communities. The ECM and microbridges as seen in our SEM images in figure 6 had a similar appearance to the fibril-like ECM material of *S*.*aureus* biofilm as seen in the A-SEM (Atmospheric-SEM) study.^59^

Multicellularity in bacteria is now a more frequently recognized behaviour^60,61^ associated with cell communication, cell-cell adhesion, biofilm formation etc.^62^ The assembly of discrete cells in a small group is considered to be the early stages of multicellularity leading to well-defined patterned clusters later. The SI_IISc_isolate with increasing concentrations of perchlorate exposure showed a greater tendency to form clusters however more experiments would be required to determine if the bacteria exhibit multicellularity which is why it is better to consider the clustering of SI_IISc_isolate to be a ‘multicellularity-like’ behaviour. In a similar study, *S*.*aureus & E*.*coli* exposed to high salinity conditions tended to grow in clusters rather than in planktonic state. A similar behaviour of environmentally induced cell-clustering was shown by *Citrobacter freundii* and *Pseudomonas aeruginosa*.^63^ This supports our idea of stress-induced multicellularity-like behaviour in SI_IISc_isolate as a response towards perchlorate.

The contribution of guar gum’s organic macromolecules towards augmentation of MICP by *Bacillus velezensis* has been established at our lab previously, where the results strongly indicated that guar gum supplementation encourages ureolysis by possibly aiding microbial survivability^51^. Our present work verifies the significant role of guar gum towards biocementation & hence continues to support the inference. Since perchlorate’s role in MICP-induced bioconsolidation of Martian regolith has been rarely explored, this study is an important insight into the Martian bricks construction via ISRU.

## V. CONCLUSION

This work is a preliminary investigation into the impact of perchlorate on (a) a lab-isolated biocementation capable bacteria phylogenetically assigned to be *Sporosarcina sp*. strain SI_IISc. (b) Martian bricks, made with MGS-1 + 1% perchlorate bioconsolidated via ureolysis-induced MICP by this strain, through an already established protocol at our lab. The results of this study suggest that perchlorate acts as a strong stress stimulus for the SI_IISc_isolate as validated by a growth curve study in broth, plate assays & live-dead staining. Multicellularity-like behaviour shown by the bacteria, observed through gram-staining images and extracellular matrix released by the bacteria observed both in gram staining & SEM images further indicated that perchlorate acted as a potent stressor for the bacteria. In SEM analysis, the extracellular matrix was observed to form microbridges, connecting the bacteria and precipitate suggesting a possible exchange between the bacteria and the precipitate.

On the other hand, the bioconsolidation experiments with Martian bricks confirmed that the biocementation capable bacteria isolated at our lab could consolidate the Martian regolith through its ureolytic activity. However, guar gum was found to be imperative for bioconsolidation, without which we could not obtain bricks of considerable compressive strength. This can be explained by guar gum’s nutritive properties for the bacteria and its role as a natural adhesive for the bricks’ consolidation. The role of NiCl_2_ as a catalyst was known through the bricks’ compressive strength which was improved as compared to the corresponding sample without nickel. In the presence of guar gum, the brick samples with perchlorate tended to have improved compressive strength as compared to the corresponding control samples without perchlorate. A possible reason for this could be the formation of inorganic precipitates by the bacteria, as observed in SEM images, which are possibly acting as binding agents. However, more studies would be needed to establish the cause. Overall, the role of various additives - guar gum, nickel, and perchlorate towards MICP-induced bioconsolidation was highlighted in the study, which underlined the complex interaction between the bacteria, regolith components, and the additives for an efficient biocementation.

