## Supplementary file for "Effect of perchlorate on biocementation capable bacteria and Martian bricks"

- **Biochemical Characterization of the SI bacteria:**

The characterization of SI bacteria was done through a series of biochemical tests using Bacillus Identification kit (HiMedia, India). The results of all biochemical tests are summarized in the table given in the table 1. The results for motility test are shown in the figure S1,

***Table S1: Summary of Biochemical tests done for SI bacteria.***

| SR. No. | Biochemical Tests | Positive/Negative |
|---|---|---|
| **1** | Malonate Utilization | N |
| **2** | Methyl Red-Vogeus Proskaeur Test or Acetoin Production | N |
| **3** | Citrate Utilization | P |
| **4** | Beta Galactosidase or ONPG test | N |
| **5** | Nitrate Utilization | N |
| **6** | Catalase | P |
| **7** | Arginine Utilization | P |
| **8** | Sucrose Utilization | N |
| **9** | Mannitol Utilization | N |
| **10** | Glucose Utilization | N |
| **11** | Arabinose Utilization | N |
| **12** | Trehalose Utilization | N |
| **13** | Urea hydrolysis | P |
| **14** | Gelatin Hydrolysis | N |
| **15** | Oxidase | N |

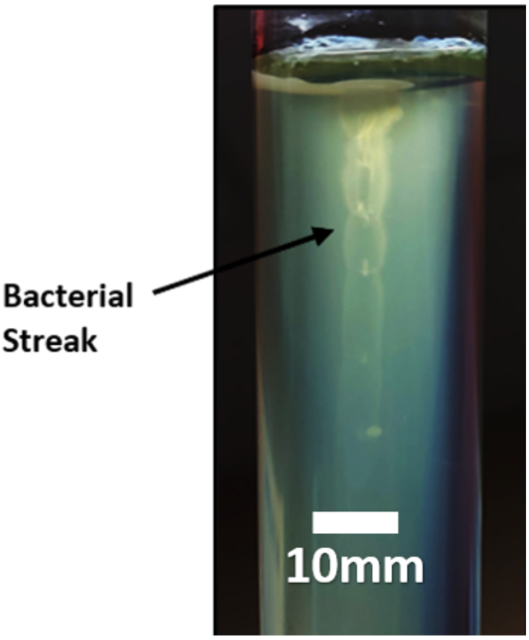


*Figure S1: Motility Test for the strain. The bacteria was introduced through an inoculation needle in Nutrient broth +2%Urea(NBU)agar media in the test tube and incubated for 48hrs at 32$^0$C. Since the bacterial streak has grown along the line of inoculation only, shows negative result for motility.*

### ➢ Ureolysis Activity in the presence of Perchlorate: Qualitative Analysis

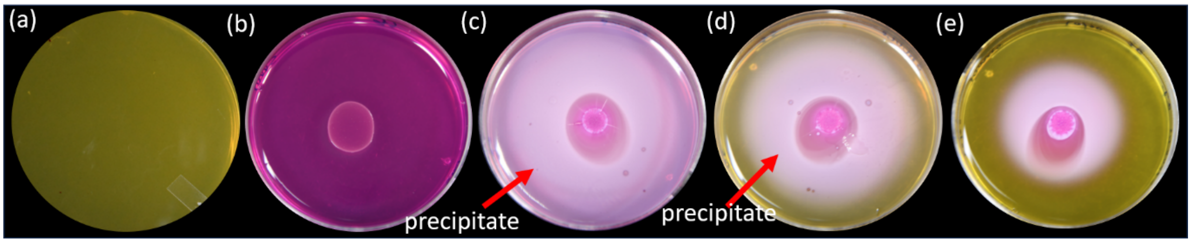


*Figure S2: Phenol red plate assay (using SMU+ 0.001%Phenol Red pH indicator dye) for the strain with varying concentrations of perchlorate (a) Control experiment with the strain (b) bacterial spot inoculation on plate with 0% perchlorate (c), (d) & (e) represents innoculation with 1%, 2% & 3% perchlorate respectively. Precipitates can be seen as white region on the plate. Ureolytic activity decreases gradually with increasing perchlorate concentration. Note: Plate dimensions- 90*14mm.*

### ➢ X-Ray Diffraction Study of microbially induced calcite precipitate

*Table S2: 2Theta values and d-spacing as matched by ICSD database (reference IDs given) through X-Ray Diffraction study of the precipitate made by SI through its ureolysis activity. The data analysis was done through X'Pert HighScore Plus Pattern Software.*

| No. | Pos. [°2Th.] | d-spacing [Å] | Rel. Int. [%] | Identified phase | hkl | ICSD Reference IDs |
|---|---|---|---|---|---|---|
| 1 | 23.12748 | 3.8427 | 9.69 | calcite | 012 | 98-004-0544 |
| 2 | 29.49984 | 3.02552 | 100 | calcite | 104 | 98-004-0544 |
| 3 | 35.99091 | 2.49335 | 10.87 | vaterite | 024 | 98-001-8127 |
| 4 | 39.45488 | 2.28206 | 16.38 | calcite | 113 | 98-004-0544 |
| 5 | 43.1715 | 2.09381 | 13.87 | vaterite | 026 | 98-001-8127 |

| | | | | | | |
|---|---|---|---|---|---|---|
| 6 | 47.62101 | 1.90804 | 15.95 | vaterite | 027 | 98-001-8127 |
| 7 | 48.57673 | 1.87271 | 15.99 | calcite | 116 | 98-004-0544 |
| 8 | 57.35916 | 1.60509 | 5.89 | calcite | 122 | 98-004-0544 |
| 9 | 60.95073 | 1.51882 | 1.62 | calcite | 208 | 98-004-0544 |
| 10 | 64.58039 | 1.44195 | 3.49 | aragonite | 122 | 98-016-1819 |

- **Effect of Perchlorate on Bacterial growth & viability**

To understand the impact of perchlorate on the bacterial growth, we determined the Minimum Inhibitory Concentration of perchlorate for the bacteria through plate assay and broth tests. To understand the effect of varying concentrations of perchlorate on the viability of our bacteria, we also performed live/dead staining using Live/Dead BacLight Kit (ThermoFischer Scientific), performed after 8 hours of incubation in nutrient media+ 2% urea (NBU)with different concentrations of perchlorate along with a control (with no perchlorate).

**(i) Minimum Inhibitory Concentration of Perchlorate**

To know the effect of perchlorate on bacteria, we first determined the minimum inhibitory concentration of perchlorate i.e. the concentration at which bacterial growth is inhibited. This was done by growth curve (quantitative)(refer Fig.S3) & plate assay (qualitative)(refer Fig.S4)

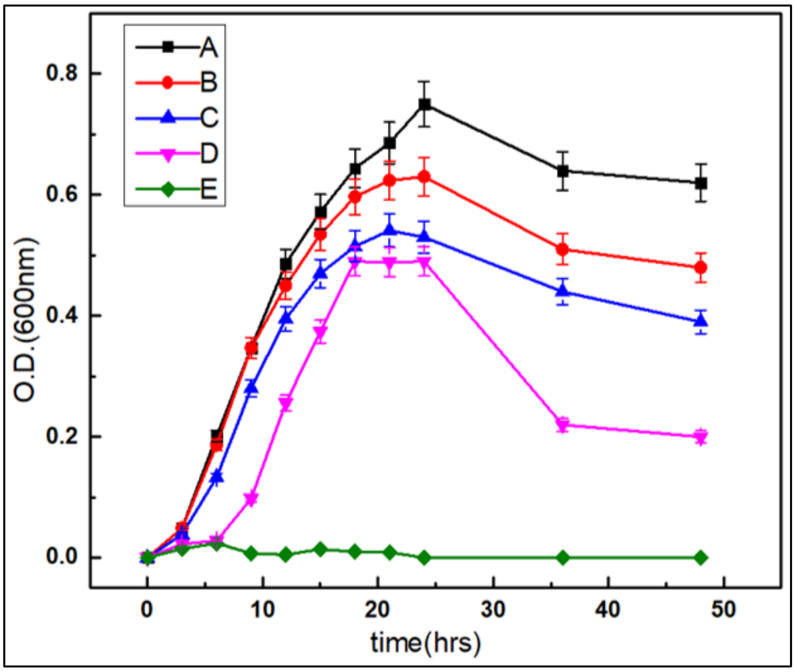


***Figure S3: Growth curve for SI_IISc bacteria in Synthetic Media + 2% Urea (SMU)broth with different concentrations of perchlorate. In the graph, A) represents growth curve for SI-(Control-No Perchlorate), b) SI+ 0.5% perchlorate C) SI +1% perchlorate D) SI+ 2%perchlorate E) SI+3% perchlorate salt. As evident from the graph, l growth of SI_IISc bacteria decreases with increasing concentration of perchlorate in the media.***

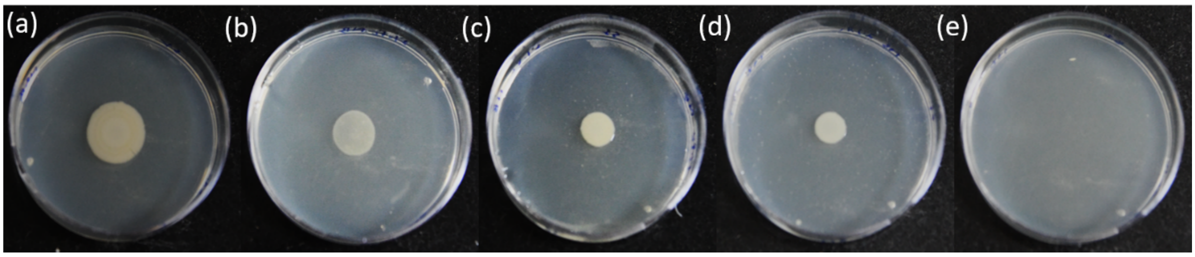


***Figure S4: Minimum Inhibitory Concentration (MIC) of Perchlorate for SI_IISc bacteria-Plate Assay- shows SI bacteria spot inoculation on Nutrient Agar plates in different concentrations of Perchlorate. (a) represents SI+ NBU (Control), 1(b) SI + 0.5% Perchlorate 1(c) SI+ 1%Perchlorate 1(d) SI+ 2%Perchlorate & (e) SI+ 3%Perchlorate. The decrease in bacterial growth is validated by decrease in diameter of bacterial spot growth, which diminishes to be zero(no growth) in (e). Note: Plate dimensions- 90*14mm.***

#### (ii) Live/Dead Staining

To understand the effect of perchlorate on viability of bacterial cells, live-dead bacterial staining was performed using Live/Dead BacLight Kit (ThermoFischer Scientific., The bacteria was grown in Nutrient Broth + 2% Urea (NBU media) & the cultures set up were (a) Control (b) SI+ 0.5% Perchlorate (c) SI + 1% Perchlorate (d) SI+2% Perchlorate.

The kit protocol was followed for the preparation of slides and were observed after 8 hours of incubation, and observed at 63X under fluorescence microscope using appropriate fluorescence filters.

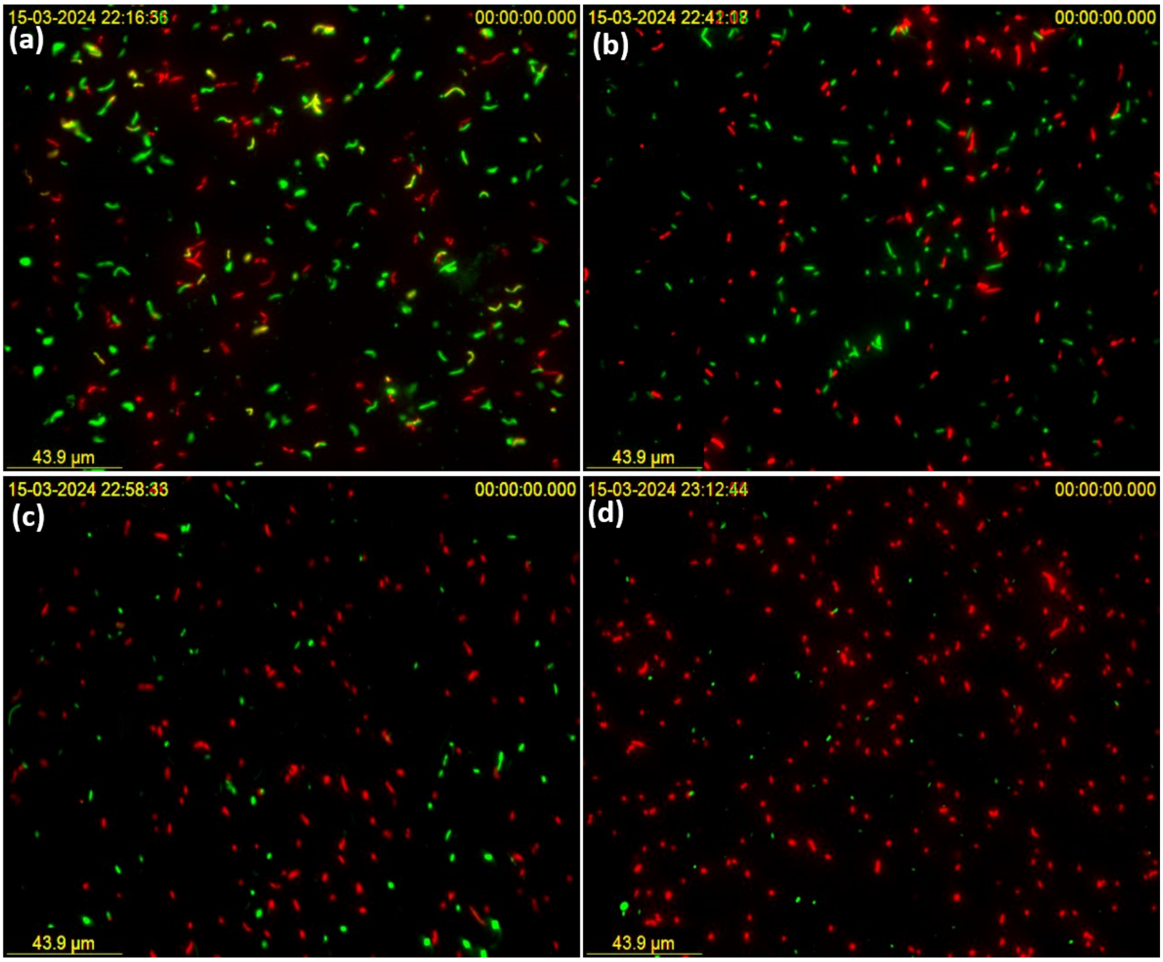


***Figure S5: Live/dead staining results after after 8 hours of incubation in Nutrient Broth +2% Urea with different concentration of perchlorate as $MgClO_4$.(a) SI(Control), (b) SI+0.5% Perchlorate (c) SI+ 1% Perchlorate (d) SI+ 2% Perchlorate. Live cells appear green while the red fluorescence***

***represent dead cells. With increasing concentration of perchlorate, the count of dead cells increased.***
